# Unexpected free fatty acid binding pocket in the cryo-EM structure of SARS-CoV-2 spike protein

**DOI:** 10.1101/2020.06.18.158584

**Authors:** Christine Toelzer, Kapil Gupta, Sathish K.N. Yadav, Ufuk Borucu, Frederic Garzoni, Oskar Staufer, Julien Capin, Joachim Spatz, Daniel Fitzgerald, Imre Berger, Christiane Schaffitzel

**Author notes:** These authors contributed equally (Christine Toelzer, Kapil Gupta, Sathish K.N. Yadav, Ufuk Borucu).

## Abstract

COVID-19, caused by severe acute respiratory syndrome-coronavirus-2 (SARS-CoV-2), represents a global crisis. Key to SARS-CoV-2 therapeutic development is unraveling the mechanisms driving high infectivity, broad tissue tropism and severe pathology. Our cryo-EM structure of SARS-CoV-2 spike (S) glycoprotein reveals that the receptor binding domains (RBDs) tightly and specifically bind the essential free fatty acid (FFA) linoleic acid (LA) in three composite binding pockets. The pocket also appears to be present in the highly pathogenic coronaviruses SARS-CoV and MERS-CoV. Lipid metabolome remodeling is a key feature of coronavirus infection, with LA at its core. LA metabolic pathways are central to inflammation, immune modulation and membrane fluidity. Our structure directly links LA and S, setting the stage for interventions targeting LA binding and metabolic remodeling by SARS-CoV-2.

**One Sentence Summary:** A direct structural link between SARS-CoV-2 spike and linoleic acid, a key molecule in inflammation, immune modulation and membrane fluidity.

## Main Text

At present, there are seven coronaviruses that are known to infect humans. The four endemic human coronaviruses OC43, 229E, HKU1 and NL63 cause mild upper respiratory tract infections while pandemic virus SARS-CoV-2, and earlier SARS-CoV and MERS-CoV, can cause severe pneumonia with acute respiratory distress syndrome, multi-organ failure, and death (*1,2*). In order to enable development of effective therapeutic interventions, a central goal of ongoing research into the COVID-19 pandemic is to determine the features of SARS-CoV-2 that provide it with a lethal combination of high infectivity and high pathogenicity.

SARS-CoV-2 has acquired novel functions that promote its harsh disease phenotype. While the previous closely related SARS-CoV did not significantly spread past the lungs, a recent study reported damage or severe inflammation in SARS-CoV-2 patients’ endothelial cells in the heart, kidneys, liver and intestines, suggestive of a vascular infection rather than a respiratory disease (*3*). The attachment of SARS-CoV-2 to a host cell is initiated by S binding to its cognate receptor angiotensin-converting enzyme 2 (ACE2), with higher affinity as compared to SARS-CoV (*4–6*). A novel S1/S2 polybasic furin protease cleavage site stimulates entry into host cells and cell-cell fusion (*4,7,8*). Inside the host cell, human coronaviruses remodel the lipid metabolism to facilitate virus replication (*9*). Infection by SARS-CoV-2 triggers an unusually impaired and dysregulated immune response (*10*) and a heightened inflammatory response (*11*) working in synergy with interferon production in the vicinity of infected cells to drive a feed-forward loop to upregulate ACE2 and further escalate infection (*12*).

In the search for additional novel functions that contribute to the observed extreme pathology of infection, we determined the structure of the SARS-CoV-2 S glycoprotein by cryo-EM (Fig.1). We discovered in our structure that the S trimer tightly binds three LA molecules in specific binding pockets, endowing a scavenger function on S, poised to impact lipid remodeling and fuel SARS-CoV-2 pathologies.

**Figure 1.**
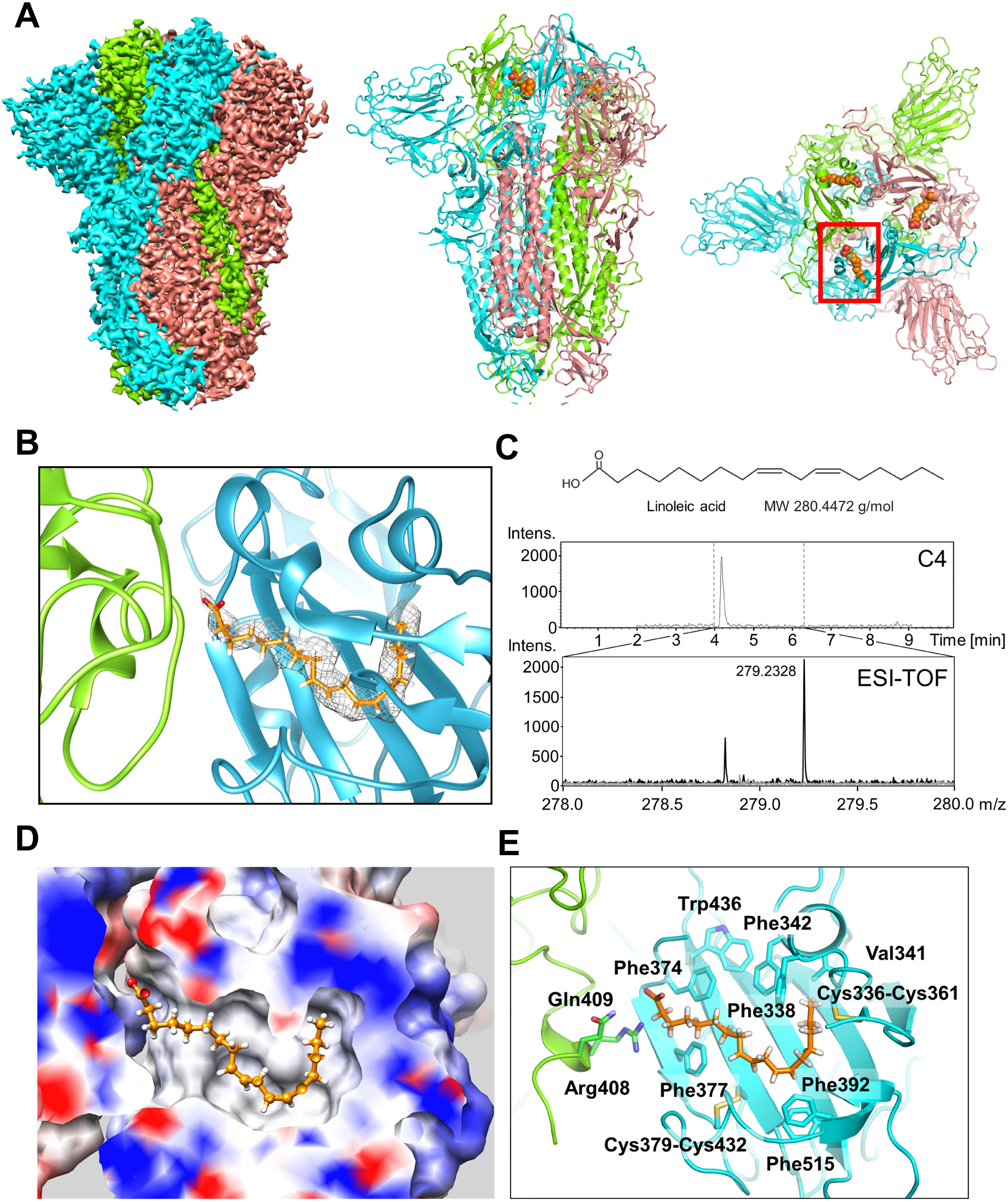
Cryo-EM structure of SARS-CoV-2 spike linoleic acid complex. (**A**) Cryo-EM density of spike trimer (left). Monomers in cyan, green and pink, respectively. The structure in a cartoon representation in a front (middle) and top view (right). Bound LA illustrated as spheres colored in orange. One LA-binding pocket is boxed in red. (**B**) Composite LA-binding pocket formed by adjacent RBDs. Tube-shaped EM density is shown. (**C**) LC-MS of purified S. Chemical structure and molecular weight of LA (top), C4 column elution profile (middle) and ESI-TOF of wash solution (grey) and C4 peak elution fraction (black) with peak molecular weight indicated (bottom). (**D**) Hydrophobic LA-binding pocket in a surface representation illustrating excellent fit of bound LA (orange, sticks and balls representation). Blue and red indicate positive and negative surface charge, respectively. (**E**) LA interactions with amino acids in the binding pocket. The acidic LA headgroup is in the vicinity of an arginine (408) and a glutamine (409).

We expressed SARS-CoV-2 S as a secreted trimer (*13*) in MultiBac (*14*) baculovirus-infected Hi5 insect cells using ESF921 media which contains cod liver oil as a nutrient supplement (Corey Jacklin, Expression Systems, personal communication). Cod liver oil comprises hundreds of FFAs including LA (*15*) while uninfected Hi5 cells contain very little LA (*16*). We purified S using immobilized metal-affinity purification and size exclusion chromatography (SEC) (fig. S1). Highly purified protein was used for cryo-EM data collection (fig. S2, table S1). After 3D classification and refinement without applying symmetry (C1) we obtained a 3.0Å closed conformation from 136,405 particles and a 3.5Å open conformation with one receptor-binding domain (RBD) in the up position from 57,990 particles (figs. S2,S3). C3 symmetry was applied to the closed conformation particle pool yielding a 2.85Å map (Fig.1A, figs. S2,S3).

The structure of S displays the characteristic overall shape observed for coronavirus S proteins in the closed and open conformations (*17–19*) with the closed form predominating in our data set (Fig.1A, figs. S2-S4). Model building of the closed form evidenced additional density in the RBDs in our structure (Fig.1B). The tube-like shape of this density was consistent with a fatty acid. Modeling conveyed LA based on size and shape similarity with LA bound to other proteins (Fig.1B, fig. S5) (*20, 21*). Liquid chromatography coupled ESI-TOF mass spectrometry (LS-MS) analysis confirmed the presence of a compound with the molecular weight of LA in our highly purified sample (Fig.1C).

Hallmarks of FFA-binding pockets in proteins are an extended ‘greasy’ tube lined by hydrophobic amino acids which accommodates the hydrocarbon tail, and a hydrophilic, often positively charged anchor for the acidic headgroup of the FFA. In our structure, a hydrophobic pocket is present which is mostly shaped by phenylalanines to form a bent tube into which the LA fits excellently (Fig.1D,E). Importantly, the anchor for the headgroup carboxyl is provided by an arginine (R408) and a glutamine (Q409) from the adjacent RBD in the trimer, giving rise to a composite LA-binding site (Fig.1E). We confirmed presence of LA in all three binding pockets in the S trimer in the unsymmetrized (C1) closed structure (fig. S6). Our S construct contains alterations as compared to native SARS-CoV-2 S including a trimerization domain and deletion of the polybasic cleavage site, neither of which alter S conformation noticeably (*13, 17*) (fig. S7). Moreover, glycosylation sites are located elsewhere and largely native in our structure (*6,17*) (table S2). We conclude that neither mutations nor glycosylation impact on the LA-binding pocket. We posit that our SARS-CoV-2 S accreted solely LA from among the many FFAs present during expression, and held on to LA firmly during purification and cryo-EM grid preparation involving massive dilutions, arguing for exquisite specificity and high affinity of the interaction.

Closed and open S conformations are in a dynamic equilibrium with the open form mediating binding to ACE2 for cell entry. We tested activity of our LA-bound S by competition enzyme-linked immunosorbent assay (ELISA) and SEC with highly purified proteins confirming ACE2 binding (Fig.2A,B). We modeled the receptor-bound structure (*22*) (Fig.2C) and found that the LA-binding pocket and the receptor binding motif (Achilles Heel, AH) of the RBD are distal and non-overlapping. The LA-bound S we produce is prolifically used in Bristol for testing patient sera for antibodies in immunology assays, further confirming functionality.

**Figure 2.**
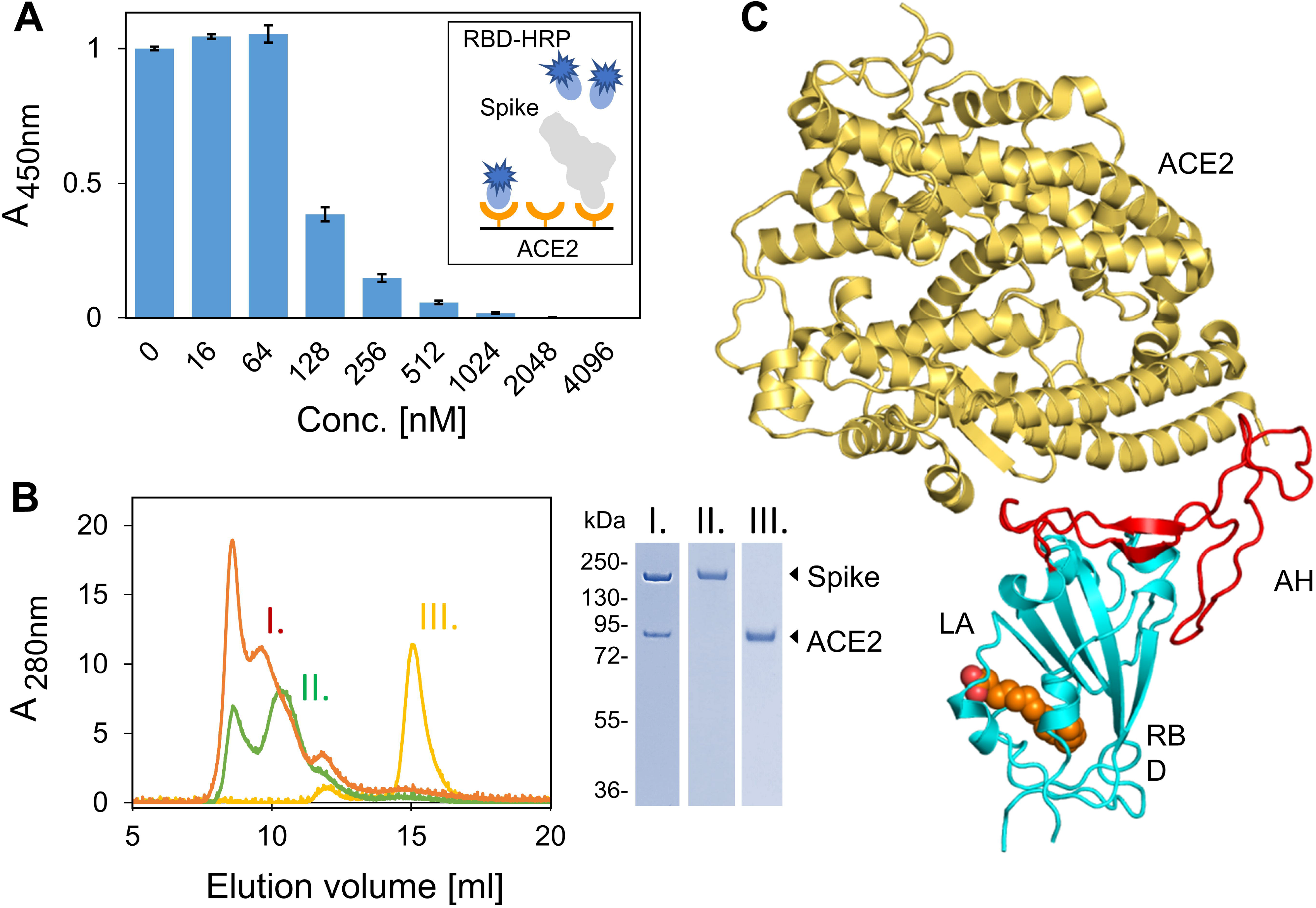
SARS-CoV-2 spike LA complex interaction with ACE2. (**A**) LA-bound SARS-CoV-2 spike displaces RBD-horse radish peroxidase (RBD-HRP) in competition ELISA utilizing immobilized ACE2. Error bars: standard deviations, three replicates. (**B**) SEC profiles for ACE2, LA-bound spike and the complex are shown. Peak fractions (I.-III.) were analysed by SDS-PAGE. (**C**) LA-bound S RBD ACE2 complex (modeled on PDBID 6M0J (*22*)). LA is shown as spheres; RBD, ACE2 in a cartoon representation. The Achilles Heel (AH, red) is fully ordered in LA-bound spike trimer in absence of ACE2, indicating conformational preorganization.

Based on our findings, we propose that the previously determined SARS-CoV-2 S structures represent an ‘apo’ form in absence of LA ligand. We hypothesize that during SARS-CoV-2 infection, S proteins of the replicating virus accrete LA giving rise to the ligand-bound form. We superimposed our LA-bound structure and the SARS-CoV-2 S ‘apo’ form (*6,17*) and identified a gating helix located directly at the entrance of the binding pocket. This gating helix, upon LA binding, moved away by about 6Å, with two tyrosines (Tyr365 and Tyr369) and Phe374 that line the gate swinging open (Fig.3A,B). In the ‘apo’ SARS-CoV-2 S trimer (*6,17*), a gap exists between adjacent RBDs with the hydrophilic anchor residues which stabilize the LA headgroup located ~10 Å away from the greasy tube entrance (Fig.3B). Upon LA binding, the gap closes, the adjacent RBD in the trimer moves towards its neighbour with anchor residues Arg408 and Gln409 locking down on the headgroup of LA (Fig.3B). Overall, this results in a marked compaction of trimer architecture in the region formed by the three RBDs (Fig.3B).

**Figure 3.**
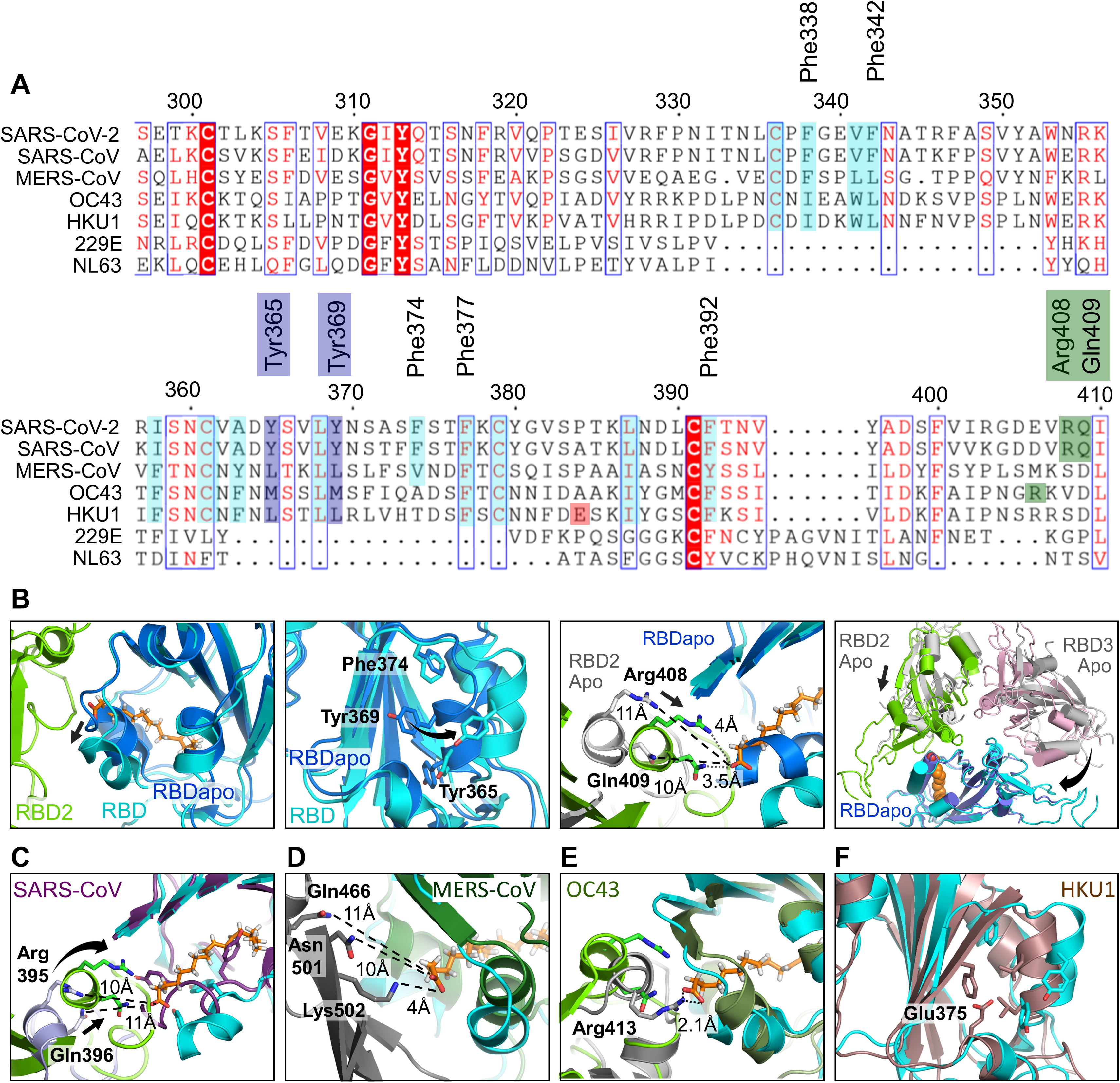
Human Coronavirus RBD architectures. (**A**) Alignments including the four common strains highlighting conserved residues. Residues lining the hydrophobic pocket are underlaid (cyan). Gating helix residues are marked (purple). Residues positioned to interact with headgroup are underlaid in green. Glutamate 375 in HKU1 is underlaid in red. (**B**) Superimposition of RBDs in LA-bound SARS-CoV-2 (cyan, green) and ligand-free ‘apo’ form (blue, grey) (PDBID 6VXX, (*6*)). Gating helix movements (left two panels), approaching of adjacent RBDs upon LA-binding (centre right) and compacting of the RBDs (far right) are illustrated. Rearrangements indicated by arrows. (**C-E**) LA-bound SARS-CoV-2 RBDs and ligand free ‘apo’ SARS-CoV (purple, PDBID 5X58 (*19*)); MERS CoV (forest green, grey, PDBID 5X5F (*19*)) and OC43 RBDs (olive, grey PDBID 6NZK, (*18*)). (**F**) HKU1 RBD (brown, PDBID 5GNB (*23*)) and RBD from LA-bound SARS-CoV-2 S (cyan).

Next, we compared the sequences and structures of human coronaviruses, including SARS-CoV-2, SARS-CoV, MERS-CoV and the four common circulating strains (Fig.3, table S3) and investigated whether the LA-binding pocket is conserved. Sequence alignment shows that all residues lining the hydrophobic pocket and the anchor residues (Arg408/Gln409) in SARS-CoV-2 are fully conserved in SARS-CoV (Fig.3A). Structural alignment of LA-bound RBDs within the trimer of SARS-CoV-2 and ‘apo’ SARS-CoV RBDs (*19*) reveals that the LA-binding pocket is present and accessible in SARS-CoV. The greasy tube is flanked by a gating helix as in SARS-CoV-2, with Arg395/Gln396 of SARS-CoV pre-positioned 10Å and 11Å from the entrance, respectively, virtually identical to apo SARS-CoV-2 (Fig.3B,C). In MERS-CoV, the gating helix and hydrophobic residues lining the pocket are also present. Tyr365, Tyr369 and Phe374 are substituted by likewise hydrophobic leucines and a valine, respectively (Fig.3A,D) (*19*). The Arg4O8/Gln4O9 pair is not conserved, however, we identify Asn501/Lys502 and Gln466 as potential anchor residues, located on a β-sheet and an α-helix within the adjacent RBD, up to 11Å away from the entrance (Fig.3D). Thus, the greasy tube and hydrophilic anchor for binding LA appear to be present in MERS-CoV, suggesting convergent evolution. In HCoV OC43, gating helix and hydrophobic residues lining the pocket are largely conserved, while Tyr365, Tyr369 and Phe374 are replaced by methionines and alanine, respectively (Fig.3A) (*18*). Arg413 is located on the same helix as Arg408/Gln409 in SARS-CoV-2 and could serve as a hydrophilic anchor (Fig.3E). No gap exists in this presumed ‘apo’ form structure between the RBDs which appear already in the locked conformation with no LA bound, likely obstructing the pocket entrance and constraining gating helix movement (Fig.3E, fig. S8) (*18*). In HCoV HKU1, the hydrophobic residues are again largely conserved, but a charged residue (Glu375) is positioned directly in front of the entrance, obstructing access for a putative LA ligand (Fig.3F) (*23*). The RBDs of HCoVs 229E and NL63, in marked contrast, adopt a very different fold (fig. S9) (*24,25*), and many of the LA-binding residues are not present (Fig.3A), indicating no obvious binding site for LA.

In summary, we find four molecular features mediating LA binding to SARS-CoV-2, and potentially also SARS-CoV and MERS-CoV S proteins: a conserved hydrophobic pocket, a gating helix, amino acid residues pre-positioned to interact with the LA carboxy headgroup, and loosely packed RBDs in the ‘apo’ form. On the other hand, in each of the four common circulating HCoVs, it appears that one or more of these four architectural prerequisites are lacking in the S protein structures (Fig. 3). LA-binding to SARS-CoV-2 S protein triggers a locking down of the hydrophilic anchor and a compaction of the RBD trimer (Fig.3B). This lockdown could contribute to the observed prevalence of closed conformation in our cryo-EM data set. It could also help stabilize the S1 region comprising the N-terminal domain and the RBD. Of note, the AH epitope, central to ACE2 binding, appears to be conformationally preorganized in our closed conformation (Fig.2C) indicating a generally more rigid RBD structure when LA is bound. While direct crosstalk in between the LA-binding pocket and the AH epitope is not apparent from our structure (Fig.2C), the overall conformational changes in the RBD trimer (Fig.3B) could conceivably impact ACE2 docking and infectivity. The S protein’s highly selective binding of LA originates from the very well-defined size and shape complementarity afforded by the LA-binding pocket (Fig. 1B,D), which is supported by the observation that we only detected LA in our analyses (Fig. 1C). The LA-binding pocket thus presents a promising target for future development of small molecule inhibitors (e.g. LA mimetics) that, for example, could irreversibly lock S in the closed conformation.

A recent proteomic and metabolomic study of COVID-19 patient sera evidenced continuous decrease of FFAs including LA (*26*). Lipid metabolome remodeling is a common element of viral infection (*27,28*), including by the baculovirus (*29*) used here to express S. For coronaviruses, the LA to arachidonic acid (AA) metabolic pathway was identified as the epicentre of lipid remodeling (*9*). Interestingly, exogenous supplement of LA or AA suppressed virus replication (*9*). It is remarkable in this context that the S trimer of SARS-CoV-2 binds LA with astonishing specificity, and we consider it highly unlikely that this is mere coincidence.

Virus-induced lipid metabolome remodeling alters three separate processes in cells (*9,27,28*): (i) energy homeostasis via changes in catabolic and anabolic precursor equilibria; (ii) fluidity and elasticity of biological membranes, via changes in e.g. saturated/unsaturated fatty acid ratio in phospholipids; and (iii) cell signaling, via changes in levels of lipid-based cell signaling precursors. By SARS-CoV-2, significant changes in cell signaling are anticipated due to LA to AA metabolome remodeling, since the LA biosynthetic pathway leads to eicosanoids, which are prominent signaling molecules involved in inflammatory processes (*30*). LA itself is detected by FFA receptor GPR40/FFAR1 (*31*). GPR40-mediated FFA signaling is a powerful mediator of inflammation in human tissues and in animal models (*32*). Regarding the potential impact of LA pathway remodeling on fluidity and elasticity of biological membranes, we note that the FFA composition of phospholipid bilayers is a key element in maintaining surface tension in lungs, and alteration of LA pathway lipid composition is observed in acute respiratory distress syndrome and severe pneumonia (*33*), both of which are key symptoms of SARS-CoV-2 infection.

The present study reveals that SARS-CoV-2 comprises a FFA-binding pocket that specifically accretes LA, and suggests that this could be a feature shared with SARS-CoV and MERS-CoV. The high affinity, high specificity LA scavenger function conveyed by our results could confer a tissue-independent mechanism by which pathogenic coronavirus infection drives immune dysregulation and inflammation. Our findings provide a direct structural link between LA, COVID-19 pathology and the virus itself and suggest that both the LA-binding pocket within the S protein and the multi-nodal LA signaling axis, represent excellent therapeutic intervention points against SARS-CoV-2 infections, particularly in patient groups with increased risk due to metabolic preconditions.

## Acknowledgments

We thank all members of the Berger and Schaffitzel teams for their contributions, Florian Krammer (Icahn School of Medicine, USA) for kindly sharing expression plasmids and Adam Finn (Children’s Vaccine Centre, Bristol Medical School), Jeremy Tavaré (School of Biochemistry, Bristol), Kathleen Gillespie (Diabetes and Metabolism Unit, Southmead Hospital, Univ. of Bristol) and Donald Fitzgerald MD (Quest Imaging Medical Associates, USA) for helpful discussions and careful reading of the manuscript. We thank Simon Burbidge, Tom Batstone and Matt Williams for computation infrastructure support. We are particularly grateful to Thiru Thangarajah (Genscript Inc.) for early access to Genscript’s cPass™ SARS-CoV-2 Neutralization Antibody Detection/Surrogate Virus Neutralization Test Kit (LOO847). We thank Sebastian Fabritz and the Core Facility for Mass Spectrometry at the Max Planck Institute for Medical Research for their support on MS measurements.

## Funding

This research received support from the Elizabeth Blackwell Institute for Health Research and the EPSRC Impact Acceleration Account EP/R511663/1, University of Bristol, from BrisSynBio a BBSRC/EPSRC Research Centre for synthetic biology at the University of Bristol (BB/LO 1386X/1) and from the BBSRC (BB/P000940/1). This work received generous support from the Oracle Higher Education and Research program to enable cryo-EM data processing using Oracle’s high-performance public cloud infrastructure (https://cloud.oracle.com/en_US/cloud-infrastructure). We acknowledge support and assistance by the Wolfson Bioimaging Facility and the GW4 Facility for High-Resolution Electron Cryo-Microscopy funded by the Wellcome Trust (202904/Z/16/Z and 206181/Z/17/Z) and BBSRC (BB/R000484/1). O.S. acknowledges support from the Elisabeth Muerer Foundation, the Max Planck School Matter to Life and the Heidelberg Biosciences International Graduate School. I.B. acknowledges support from the EPSRC Future Vaccine Manufacturing and Research Hub (EP/R013764/1). C.S. and I.B. are Investigators of the Wellcome Trust (210701/Z/18/Z; 106115/Z/14/Z).

## Author Contributions

C.S. and I.B. conceived and guided the study. F.G. and K.G. produced and purified sample, K.G. carried out biochemical experiments, S.K.N.Y. and U.B. prepared grids and collected EM data, S.K.N.Y., U.B., K.G and C.T. carried out image analysis and model building. O.S. and J.S. performed and interpreted mass spectrometry. J.C. prepared reagents and analysed data. C.T., K.G., D.F., I.B. and C.S. interpreted data. D.F., I.B. and C.S. wrote the manuscript with input from all authors.

## Competing interests

The authors declare competing interests. I.B. and D.F. report shareholding in Geneva Biotech Sàrl unrelated to this Correspondence. I.B and F.G. report shareholding in Imophoron Ltd. unrelated to this Correspondence. A patent application describing drug discovery methods and therapeutic interventions based on the present observations has been filed.

## Data and materials availability

Datasets generated during the current study have been deposited in the Electron Microscopy Data Bank (EMDB) under accession numbers EMD-11145 (C3 closed conformation), EMD-11144 (C1 closed conformation) and EMD-11146 (open conformation), and in the Protein Data Bank (PDB) under accession numbers: 6ZB5 (C3 closed conformation) and 6ZB4 (C1 closed conformation).

## Notes

### Competing Interest Statement

I.B. and D.F. report shareholding in Geneva Biotech SARL unrelated to this Correspondence. I.B and F.G. report shareholding in Imophoron Ltd. unrelated to this Correspondence. A patent application describing drug discovery methods and therapeutic interventions based on the present observations has been filed.

