## Supplementary Materials for "Unexpected free fatty acid binding pocket in the cryo-EM structure of SARS-CoV-2 spike protein"

### Supplement

### Materials and Methods

Protein production.

*SARS-CoV-2 spike protein.* The pFastBacDual plasmid for SARS-CoV-2 S ectodomain for expression in insect cells (*13*) was kindly provided by Florian Krammer (Icahn School of Medicine, USA). In this construct, S comprises amino acids 1 to 1213 and is fused to a C-terminal thrombin cleavage site followed by a T4-foldon trimerization domain and a hexahistidine affinity purification tag. The polybasic cleavage site has been deleted (RRAR to A) in the construct (*13*). Protein was produced with the MultiBac baculovirus expression system (Geneva Biotech, Geneva, Switzerland) (*34*) in Hi5 cells using ESF921 media (Expression Systems Inc.). Supernatants from transfected cells were harvested 3 days post-transfection by centrifugation of the culture at 1,000g for 10 min followed by another centrifugation of supernatant at 5,000g for 30 min. The final supernatant was incubated with 7 ml HisPur Ni-NTA Superflow Agarose (Thermo Fisher Scientific) per 3 litres of culture for 1h at 4ºC. Subsequently, a gravity flow column was used to collect the resin bound with SARS-CoV-2 spike protein, the resin was washed extensively with wash buffer (65 mM $NaH_2PO_4$, 300 mM NaCl, 20 mM imidazole, pH 7.5), and the protein was eluted using a step gradient of elution buffer (65 mM $NaH_2PO_4$, 300mM NaCl, 235mM imidazole, pH 7.5). Elution fractions were analysed by reducing SDS-PAGE and fractions containing SARS-CoV-2 spike protein were pooled, concentrated using 50 kDa MWCO Amicon centrifugal filter units (EMD Millipore) and buffer-exchanged in phosphate-buffered saline (PBS) pH 7.5. Concentrated SARS-CoV-2 S was subjected to size exclusion chromatography (SEC) using a Superdex 200 increase 10/300 column (GE Healthcare) in PBS pH 7.5. Peak fractions from SEC were analysed by reducing SDS-PAGE and negative stain electron microscopy (EM); fraction 7 was used for cryo-EM (fig.S1).

*ACE2 protein.* The gene encoding for the ACE2 ectodomain (amino acids 1 to 597, (*35*)) was codon optimized for insect cell expression, synthesized (Genscript Inc, New Jersey USA) and inserted into pACEBac1 plasmid (Geneva Biotech, Geneva, Switzerland). The construct contains an N-terminal melittin signal sequence for secretion and a C-terminal octahistidine affinity purification tag. ACE2 protein was produced and purified following the same protocol as for spike protein.

Negative stain sample preparation and microscopy

4 μL of 0.05 mg/mL SARS-CoV-2 spike protein was applied onto a freshly glow discharged (1 min at 10 mA) CF300-Cu-50 grid (Electron Microscopy Sciences), incubated for 1 min, and manually blotted. 4 μL of 3% Uranyl Acetate was applied onto the same grid and incubated for 1 min before the solution was blotted off. The grid was loaded onto a FEI Tecnai 12 120 kV BioTwin Spirit TEM. Images were acquired at a nominal magnification of 49,000x.

Cryo-EM sample preparation and data collection

4 μL of 1.25 mg/mL SARS-CoV-2 spike protein was loaded onto a freshly glow discharged (2 min at 4 mA) Quantifoil R1.2/1.3 carbon grid (Agar Scientific), blotted using a Vitrobot MarkIV (Thermo Fisher Scientific) at 100% humidity and 4°C for 2 s, and plunge frozen. Data were acquired on a FEI Talos Arctica transmission electron microscope operated at 200 kV and

equipped with a Gatan K2 Summit direct detector and Gatan Quantum GIF energy filter, operated in zero-loss mode with a slit width of 20 eV using the EPU software.

Data were collected in super-resolution at a nominal magnification of 130,000x with a virtual pixel size of 0.525 Å. The dose rate was adjusted to 6.1 counts/physical pixel/s. Each movie was fractionated in 55 frames of 200 ms. 3289 micrographs were collected in a single session with a defocus range comprised between -0.8 and -2 μm.

Cryo-EM data processing

The dose-fractionated movies were gain-normalised, aligned, and dose-weighted using MotionCor2 (*36*). Defocus values were estimated and corrected using CTFFIND4 (*37*). 611,879 particles were automatically picked using Relion 3.0 software (*38*). Reference-free 2D classification was performed to select well-defined particles, after four rounds of 2D classification a total of 386,510 good particles were selected for further 3D classification. An initial model was generated using 50,000 particle images in Relion 3.0, and the selected particles from 2D classification were subjected to 3D classification using 8 classes. Classes 4 and 6 (fig. S2), showing prominent features representing a total of 202,082 particles were combined and used for 3D refinement. The 3D refined particles were then subjected to a second round of 3D classification using 5 classes. Class 4 and 5 were combined for the closed conformation map, class 3 represented the open conformation, comprising 136,405 and 57,990 particles respectively. The selected maps were subjected to 3D refinement without applying any symmetry with respective 3D models. The maps were subsequently subjected to local defocus correction and Bayesian particle polishing in Relion 3.0. Global resolution and B factor (-89Å$^2$ and -116Å$^2$ for closed and open maps respectively) of the map were estimated by applying a soft mask around the protein density, using the gold standard Fourier shell correlation (FSC) = 0.143 criterion, resulting in an overall resolution of 3.03Å and 3.7Å respectively. The open map was imported into Relion 3.1 software, where it was further corrected for higher-order aberrations and magnification anisotropy. This yielded a final resolution of 3.5 Å (B factor of -87$Å^2$). C3 symmetry was applied to the closed conformation map using Relion 3.0, followed by 3D classification using 3 classes. Class 3 with 217,815 particles was selected for CTF refinement and Bayesian polishing, yielding a final resolution of 2.85Å (B factor of -86.8). Local resolution maps were generated using Relion 3.0 (fig. S3).

Cryo-EM model building and analysis

UCSF Chimera (*39*) and Coot (*40*) were used to fit atomic models (PDBID 6VXX, (*6*)) into the cryo-EM map. The model was subsequently manually rebuilt using Coot and the closed conformation map. This closed conformation model was used to build features specific of the open conformation map and to further improve the model by using the C3-symmetrized map. To guide the model building, sharpened (*41*) and unsharpened maps were used in different steps of the process. N-linked glycans were hand-built into the density where visible, and restraints for non-standard ligands were generated with eLBOW (*42*). The model for the closed conformation was real space refined with Phenix (*43*), and the quality was additionally analysed using MolProbity (*44*) and EMRinger (*45*), to validate the stereochemistry of the components. The quasi-atomic model of the open conformation was generated by first fitting the atomic model of the closed conformation into the open cryo-EM map using UCSF Chimera. We then used COOT and UCSF Chimera to move one RBD into the open conformation, by aligning this RBD to the atomic model of a published open form of SARS-CoV-2 S (PDBID 6VSB, (*17*)). Finally, this open model was fitted into the cryo-EM open map with COOT and UCSF Chimera. Figures were prepared using UCSF chimera and PyMOL (Schrodinger, Inc).

Mass spectrometry analysis

For mass spectrometry analysis, a Bruker maXis II ETD quadrupole-time-of-flight instrument with electrospray ionization (ESI) coupled to a Shimadzu Nexera HPCL was used. In order to extract the fatty acids from the protein sample, a chloroform extraction protocol was performed. For this, 100 µl of the protein sample was mixed with 400 µl chloroform for 2 hours on a horizontal shaker in a teflon-sealed glass vial at 25°C. The organic phase was then transferred to a new glass vial and the chloroform was evaporated for 30 min in a desiccator. Subsequently, 50 µl of a 20% (v/v) acetonitrile solution was added to dissolve the fatty acids. From this solution, a 1:100 dilution in 20% (v/v) acetonitrile was prepared and 20 µl were injected for LC-MS analysis. The samples were passed over a Phenomenex C4 Aeris column (100 × 2.1 mm, 3.6µ, 200 Å) heated to 50°C using a gradient of 20% A to 98% B in 5 min (solvent A: $H_2O$ + 0.1% formic acid; solvent B: ACN + 0.1% formic acid). The system was operated with Bruker's O-TOF Control (V4.1) and Hystar (V4.1) software and data analysis as well as data post-processing was performed with Bruker's DataAnalysis software (V4.4, SR1).

ELISA activity assay

The SARS-CoV-2 sVNT kit was obtained from GenScript Inc. (New Jersey, USA) for ELISA activity assays. Serial dilution series were prepared (from 0-4096 nM) of purified SARS-CoV-2 spike protein in PBS pH 7.5. This dilutions series was mixed with the same amount of HRP-RBD and incubated at 37ºC for 30 min. The mixture of each dilution of SARS-CoV-2 spike proteins and HRP-RBD was then added to ELISA plate wells, which were coated with ACE2. Triplicates of each sample were added. The ELISA plate was then incubated at 37ºC for 15 min and subsequently washed 4 times with wash buffer. The signal was then developed by adding TMB solution to each well, incubating 15 minutes in the dark, followed by adding stop solution. The absorbance at 450nm was immediately recorded. The data was plotted using Microsoft excel. The standard deviation of triplicates was added as error bars.

<u>SARS-CoV-2 spike and ACE2 interaction analysis</u>

Purified SARS-CoV-2 spike and ACE2 proteins were combined in PBS pH 7.5 at a ratio of 1:1.5 spike trimer to ACE2 monomer. The mixture was incubated on ice for 2 hours and subjected to SEC using a Superdex 200 increase 10/300column (GE Healthcare) equilibrated in PBS pH 7.5. Purified SARS-CoV-2 spike protein and ACE2 proteins were individually run on the same column in the same buffer as controls. The contents of peak fractions were analysed by reducing SDS-PAGE.

**Figure S1.**

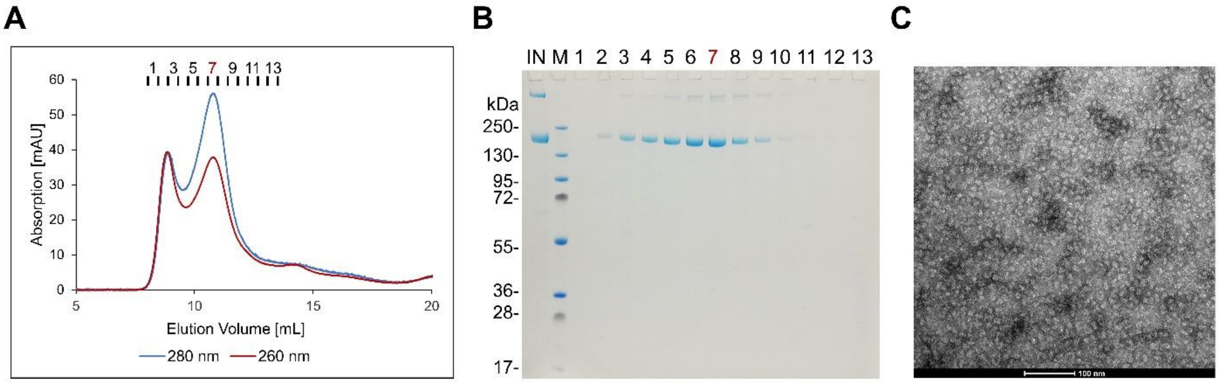


**Purification and quality control of SARS-CoV-2 spike protein**. (**A**) Size-exclusion chromatogram of affinity-purified SARS-CoV-2 spike protein using a Superdex 200 column. Absorption was detected at 280 nm (blue line) and 260 nm (red line). Peak fractions are indicated. (**B**) SDS PAGE analysis of the SEC fractions from panel A. Lane 1: input fraction, lane 2: molecular weight marker, lane 3-15: fractions 1 to 13 from SEC. (**C**) Negative stain EM of SEC fraction 7 (scale bar: 100 nm).

**Figure S2.**

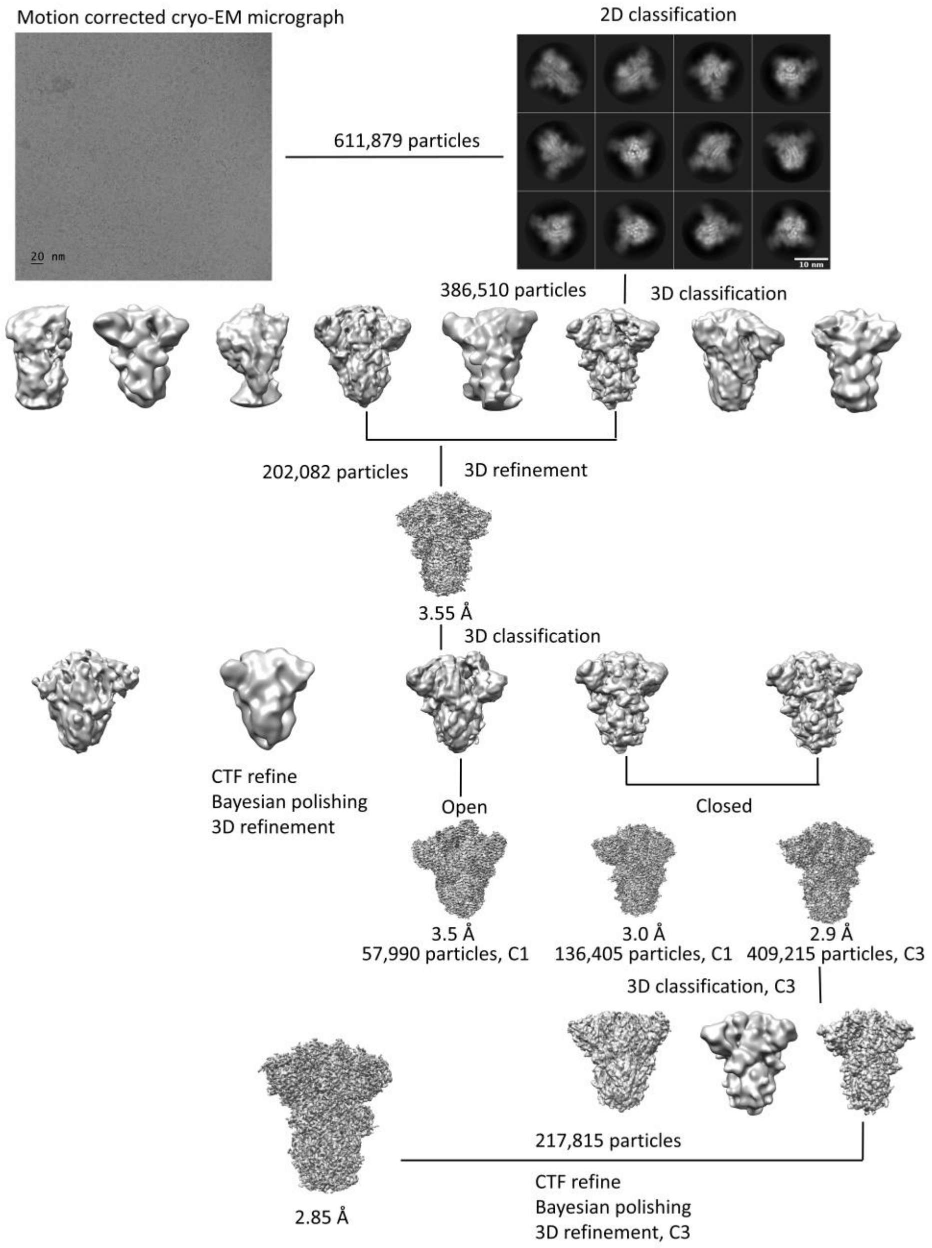


**Cryo-EM image processing workflow**. A motion-corrected cryo-EM micrograph is shown (scale bar 20 nm), reference-free 2D class averages (scale bar 10 nm), 3D classification and refinement resulting in cryo-EM maps corresponding to the open conformation and the closed conformation (not symmetrized and C3 symmetrized).

**Figure S3.**

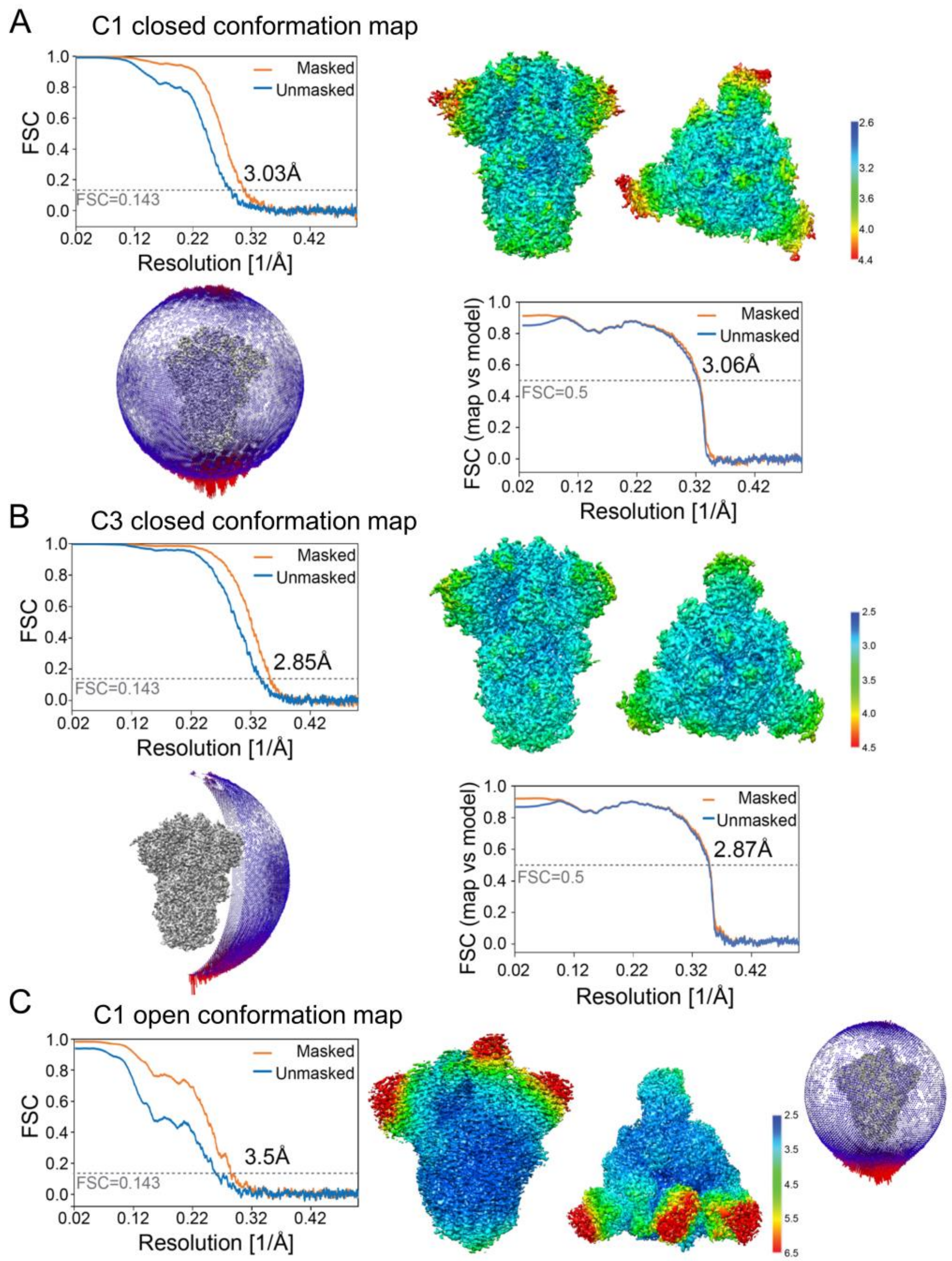


**Cryo-EM structure validation.** Top left: Fourier Shell Correlation (FSC) curve after gold standard refinement. Top right: Cryo-EM reconstruction colored according to the local resolution from a side and top view. Below left: Orientation distribution of views that contributed to this map.

Longer red rods represent orientations that comprise more particles. Below right: Cross-validation FSC curves for the refined model versus the final masked and unmasked maps, shown for (**A**) the closed unsymmetrized C1 map (**B**) the closed C3-symmetrized map and (**C**) the open conformation map. For the open conformation, the views distribution is shown on the right, next to the local

**Figure S4.**

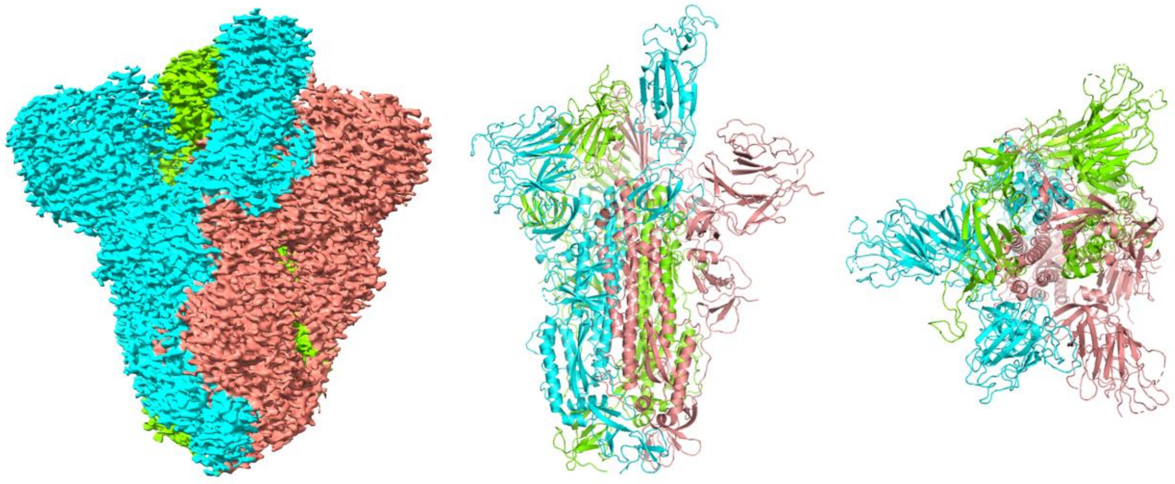

**Cryo-EM structure of SARS-CoV-2 S in the open conformation**. The 3.5Å cryo-EM density of the SARS-CoV-2 spike trimer is shown (left). Monomers are colored in cyan, green and pink, respectively. A quasi-atomic model is depicted in a cartoon representation in a front (middle) and top view (right) using the same colors (c.f. Fig.1). RBDs were placed as rigid bodies, and LA was not included in the model.

**Figure S5.**

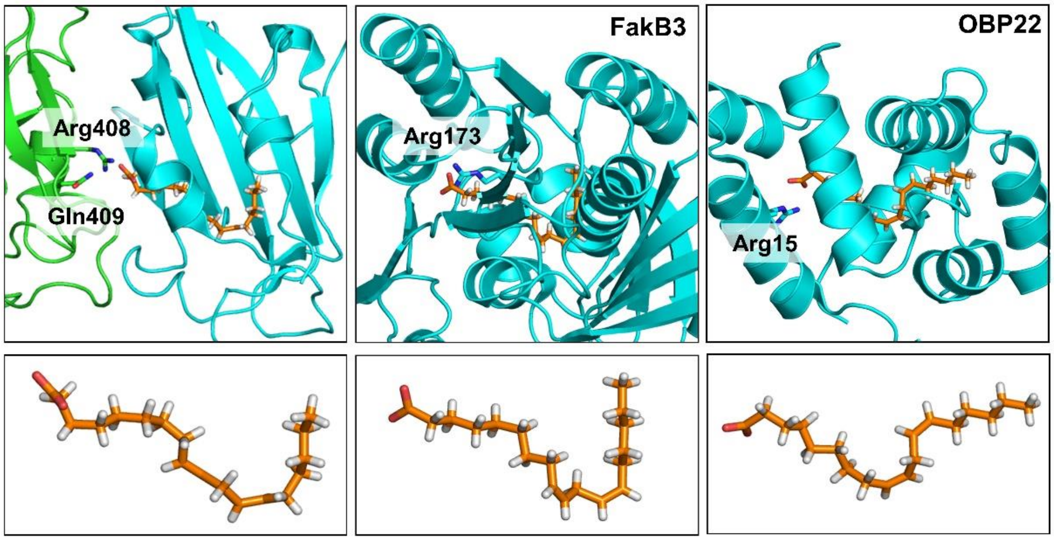


**LA binding protein complexes**. The structures of LA bound to SARS-CoV-2 S protein (this study, left), *Streptococcus pneumoniae* Fatty Acid Kinase (Fak) B3 (PDBID 6CNG (*20*), middle) and *Aedes aegypti* Odorant Binding Protein 22 (OBP22) (PDBID 6OGH, (*21*), right) are shown for comparison (top panel). Amino acids interacting with the polar headgroup of LA are labelled. Despite markedly different protein architectures, the conformation adopted by the LA in each complex is similar, featuring a pronounced kink in the structures (close-up views below).

**Figure S6.**

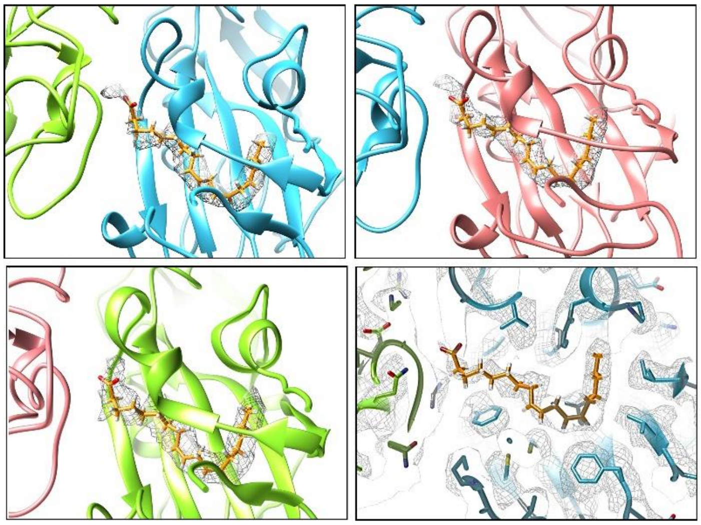

**LA ligands bound to the SARS-CoV-2 spike RBDs**. The three independent RBDs present in SARS-CoV-2 spike trimer are shown. Processing and refinement in C1 (no symmetry applied) results in tube-shaped density within all three RBDs, indicating that a LA ligand is present in each fatty acid-binding pocket. EM density is depicted as a mesh colored in grey. RBDs are colored cyan, green and pink (c.f. Figure 1). LA ligand is shown as sticks colored in orange.  The density of amino acid residues in the binding pocket surrounding LA in the C3 structure is shown for comparison in a clipped view (bottom right).

**Figure S7.**

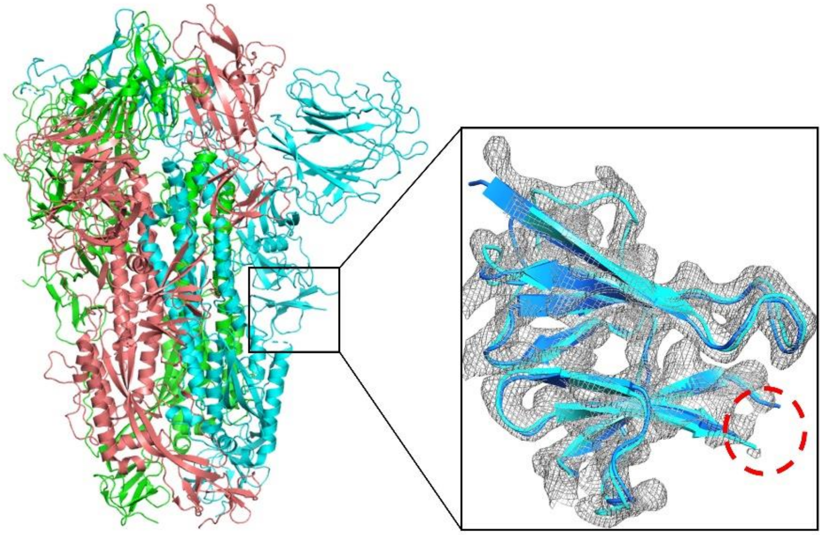

**Polybasic cleavage site deletion in SARS-CoV-2 spike.** The construct used in this study has a RRAR to A deletion in the polybasic cleavage site (boxed in black). This site is part of a flexible loop in the S protein which is not observed by cryo-EM (circled in red in close-up view). Comparison of the LA-bound SARS-CoV-2 spike protein (cyan) and the 'apo' form (dark blue, PDBID 6VXX, RRAR->SGAG (*6*)) shown in a close up view superimposed on cryo-EM density (grey mesh) evidences that the structures are virtually identical in this region.

**Figure S8.**

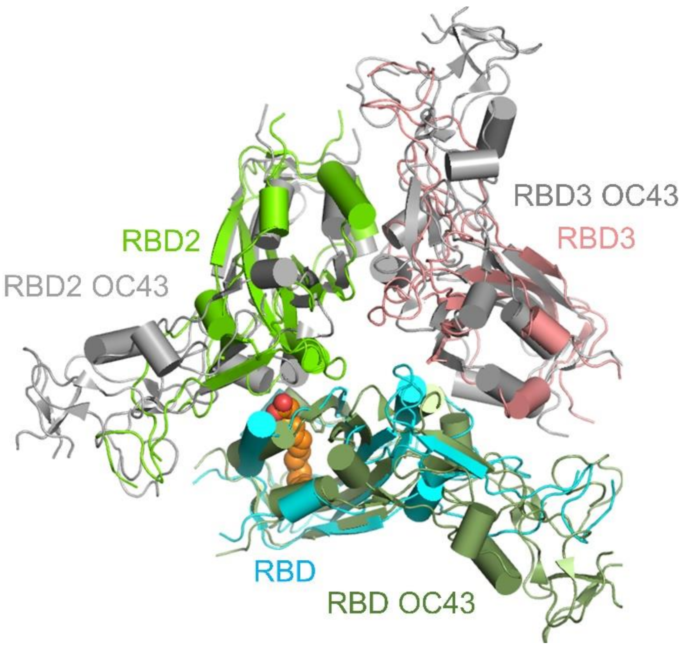


**Superimposition of RBDs in OC43 S protein trimer and LA-bound SARS-CoV-2 S.** The OC43 RBD colored in dark green and the LA-bound SARS-CoV-2 RBD colored in cyan were used for structure alignment. Two further RBDs in the trimer in OC43 S are colored in grey. Corresponding RBDs in the LA-bound SARS-CoV-2 S are colored in green and pink (c.f. Fig. 1). The three RBDs of OC43 are locked in a tightly packed conformation with close contacts at the RBD-RBD interfaces (PDBID 6NZK (*18*)).

**Figure S9.**

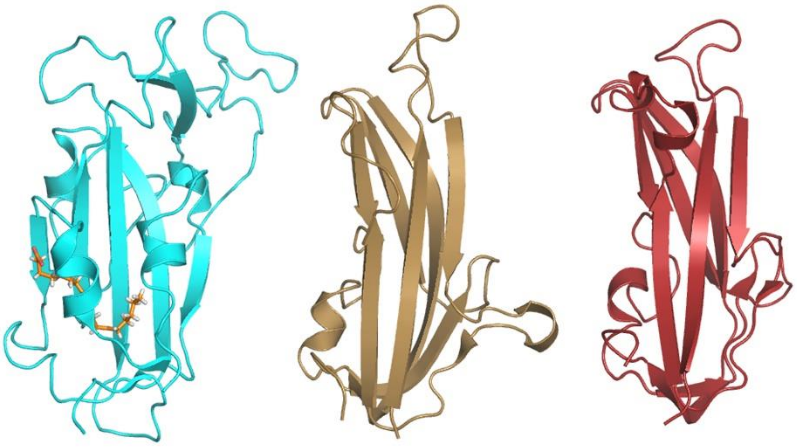


**Spike protein RBD structures**. LA-bound SARS-CoV-2 S RBD (cyan, this study, left) and RBDs of commonly circulating human coronaviruses 229E (PDBID 6U7H (*24*), middle) and NL63 (PDBID 5SZS (*25*), right) are shown. The RBD architectures of 229E and BL63 are substantially different.

**Table S1.**

**Cryo-EM data collection and refinement statistics.**

| | **Closed conformation, C3 symmetrized** | **Closed conformation, C1** | **Open conformation, C1** |
|---|---|---|---|
| Voltage (kV) | 200 | 200 | 200 |
| Magnification (nominal) | 130,000 | 130,000 | 130,000 |
| Pixel size (Å/pix) | 1.05 (0.525) | 1.05 (0.525) | 1.05 (0.525) |
| Flux ($e^-$ /pix/sec) | 6.1 | 6.1 | 6.1 |
| Frames per exposure | 55 | 55 | 55 |
| Exposure ($e^-$ /Å$^2$) | 60.5 | 60.5 | 60.5 |
| Defocus range (mm) | -0.8 to -2 | -0.8 to -2 | -0.8 to -2 |
| Micrographs collected | 3289 | 3289 | 3289 |
| Particles final | 217,815 | 136,405 | 57,990 |
| Map sharpening B-factor (Å$^2$) | -86.8 | -89 | -87 |
| Masked resolution at 0.143 FSC (Å) | 2.85 | 3.03 | 3.5 |

**Refinement**

| | **Closed conformation, C3 symmetrized** | **Closed conformation, C1** |
|---|---|---|
| Composition | | |
| Amino acids | 3127 | 3133 |
| Glycans | 27 | 27 |
| Ligands | 3 | 3 |
| RMSD bonds (Å) | 0.003 | 0.002 |
| RMSD angles (º) | 0.502 | 0.501 |
| Mean B-factors (Å$^2$) | | |
| Amino acids | 21.88 | 30.13 |
| Ligands | 42.56 | 46.99 |
| Ramachandran | | |
| Favoured (%) | 92.60 | 91.32 |
| Allowed (%) | 7.37 | 8.59 |
| Outliers (%) | 0.03 | 0.10 |
| Rotamer outliers (%) | 5.49 | 4.35 |
| Clash score | 9.42 | 14.90 |
| C-beta outliers (%) | 0.00 | 0.00 |
| CaBLAM outliers (%) | 3.56 | 4.14 |
| CC (mask) | 0.86 | 0.84 |
| MolProbity score | 2.52 | 2.67 |
| EMRinger score | 3.43 | 3.64 |
| Model resolution (Å) 0.5 FSC threshold | 2.9 | 3.1 |

**Table S2.**
**N-linked glycosylation sites in the SARS-CoV-2 spike Protein**

| WT SARS-CoV-2 spike * | Recombinant SARS-CoV-2 spike expressed in | |
|---|---|---|
| **Predicted (UNIPROT)** | **Freestyle 293F (*6*) ** ** | **Hi5 (this study) ** ** |
| $N_{17}$LT | | $N_{17}$LT |
| $N_{61}$VT | $N_{61}$VT | $N_{61}$VT |
| $N_{74}$GT | | |
| $N_{121}$NA [#] | | |
| $N_{122}$AT | $N_{122}$AT | $N_{122}$AT |
| $N_{149}$KS | | |
| $N_{165}$CT | $N_{165}$CT | $N_{165}$CT |
| $N_{234}$IT | $N_{234}$IT | $N_{234}$IT |
| $N_{282}$GT | $N_{282}$GT | $N_{282}$GT |
| $N_{331}$IT | $N_{331}$IT | $N_{331}$IT |
| $N_{343}$AT | $N_{343}$AT | $N_{343}$AT |
| $N_{370}$SA [#] | | |
| $N_{603}$TS | $N_{603}$TS | |
| $N_{616}$CT | $N_{616}$CT | $N_{616}$CT |
| $N_{657}$NS | $N_{657}$NS | |
| $N_{709}$NS | $N_{709}$NS | $N_{706}$NS |
| $N_{717}$FT | $N_{717}$FT | $N_{714}$FT |
| $N_{801}$FS | $N_{801}$FS | $N_{798}$FS |
| $N_{1074}$FT | $N_{1074}$FT | $N_{1071}$FT |
| $N_{1098}$GT | $N_{1098}$GT | $N_{1095}$GT |
| $N_{1134}$NT | $N_{1134}$NT | $N_{1131}$NT |
| $N_{1158}$HT | | |
| $N_{1173}$AS | | |
| $N_{1194}$ES | | |

***** *YP_009724390.1*
***Sites lacking glycosylation in cryo-EM maps are omitted (boxes colored in grey).*
[#] *Predicted based on SARS-CoV spike protein (6)*

**Table S3.**

**Alignment of coronavirus S protein sequences** (S-CryoEM denotes construct in this study)

```
                                        940       950       960       970
S-CryoEM     IQDSLSSTA..............SALGKLQDVVNQNAQALNTLVKQLSSNFGAISSVLND
SARS-CoV-2   IQDSLSSTA..............SALGKLQDVVNQNAQALNTLVKQLSSNFGAISSVLND
SARS-CoV     IQESLTTTS..............TALGKLQDVVNQNAQALNTLVKQLSSNFGAISSVLND
MERS-CoV     MQTGFTTTN..............EAFRKVQDAVNNNAQALSKLASELSNTFGAISASIGD
HCoV-229E    IVDAFTGVNDAITQTSQALQTVATALNKIQDVVNQQGNSLNHLTSQLRQNFQAISSSIQA
HCoV-OC43    IQQGFDATN..............SALVKIQAVVNANAEALNNLLQQLSNRFGAISASLQE
HCoV-NL63    IVASFSSVNDAITQTAEAIHTVTIALNKIQDVVNQQGSALNHLTSQLRHNFQAISNSIQA
HCoV-HKU1    IQNGFSATN..............SALAKIQSVVNSNAQALNSLLQQLFNKFGAISSSLQE

               980       990       1000      1010      1020      1030
S-CryoEM     ILSRLDKVEAEVQIDRLITGRLQSLQTYVTQQLIRAAEIRASANLAATKMSECVLGQSKR
SARS-CoV-2   ILSRLDKVEAEVQIDRLITGRLQSLQTYVTQQLIRAAEIRASANLAATKMSECVLGQSKR
SARS-CoV     ILSRLDKVEAEVQIDRLITGRLQSLQTYVTQQLIRAAEIRASANLAATKMSECVLGQSKR
MERS-CoV     IIQRLDVLEQDAQIDRLINGRLTTLNAFVAQQLVRSESAALSAQLAKDKVNECVKAQSKR
HCoV-229E    IYDRLDTIQADQQVDRLITGRLAALNVFVSHTLTKYTEVRASRQLAQQKVNECVKSQSKR
HCoV-OC43    ILSRLDALEAEAQIDRLINGRLTALNAYVSQQLSDSTLVKFSAAQAMEKVNECVKSQSSR
HCoV-NL63    IYDRLDSIQADQQVDRLITGRLAALNAFVSQVLNKYTEVRGSRRLAQQKINECVKSQSNR
HCoV-HKU1    ILSRLDALEAQVQIDRLINGRLTALNAYVSQQLSDISLVKFGAALAMEKVNECVKSQSPR

             1040      1050      1060      1070      1080         1090
S-CryoEM     VDFCGKGYHLMSFPQSAPHGVVFLHVTYVPAQEKNFTTAPAICHDGKA...HFPREGVFV
SARS-CoV-2   VDFCGKGYHLMSFPQSAPHGVVFLHVTYVPAQEKNFTTAPAICHDGKA...HFPREGVFV
SARS-CoV     VDFCGKGYHLMSFPQAAPHGVVFLHVTYVPSQERNFTTAPAICHEGKA...YFPREGVFV
MERS-CoV     SGFCGQGTHIVSFVVNAPNGLYFMHVGYYPSNHIEVVSAYGLCDAANPTNCIAPVNGYFI
HCoV-229E    YGFCGNGTHIFSIVNAAPEGLVFLHTVLLPTQYKDVEAWSGLCVDGTN..GYVLRQPNLA
HCoV-OC43    INFCGNGNHIISLVQNAPYGLYFIHFNYVPTKYVTAKVSPGLCIAGNR..GIAPKSGYFV
HCoV-NL63    YGFCGNGTHIFSIVNSAPDGLLFLHTVLLPTDYKNVKAWSGICVDGIY..GYVLRQPNLV
HCoV-HKU1    INFCGNGNHILSLVQNAPYGLLFMHFSYKPISFKTVLVSPGLCISGDV..GIAPKQGYFI

                       1100      1110      1120      1130        1140
S-CryoEM     SN.....GTHWFVTQRNFYEPQIITTDNTFVSGNCDVVIGIVNNTVYDPLQ..PELDSFK
SARS-CoV-2   SN.....GTHWFVTQRNFYEPQIITTDNTFVSGNCDVVIGIVNNTVYDPLQ..PELDSFK
SARS-CoV     FN.....GTSWFITQRNFFSPQIITTDNTFVSGNCDVVIGIINNTVYDPLQ..PELDSFK
MERS-CoV     KTNNTRIVDEWSYTGSSFYAPEPITSLNTKYVAPQVTYQN.ISTNLPPPLLGNSTGIDFQ
HCoV-229E    LY...KEGNYYRITSRIMFEPRIPTMADFVQIENCNVTFVNISRSEL..QTIVPEYIDVN
HCoV-OC43    NV.....NNTWMYTGSGYYYPEPITENNVVVMSTCAVNYTKAPYVML..NTSIPNLPDFK
HCoV-NL63    LY...SDNGVFRVTSRVMFQPRLPVLSDFVQIYNCNVTFVNISRVEL..HTVIPDYVDVN
HCoV-HKU1    KH.....NDHWMFTGSSYYYPEPISDKNVVVFMNTCSVNFTKAPLVYL..NHSVPKLSDFE

             1150        1160        1170                  1180     1190
S-CryoEM     EELDKYFKNHTS.PDVDLG.DISGINASVVNI..............QKEIDRLNEVAKNL
SARS-CoV-2   EELDKYFKNHTS.PDVDLG.DISGINASVVNI..............QKEIDRLNEVAKNL
SARS-CoV     EELDKYFKNHTS.PDVDLG.DISGINASVVNI..............QKEIDRLNEVAKNL
MERS-CoV     DELDEFFKNVST.SIPNFG.SLTQINTTLLDL..............TYEMLSLQQVVKAL
HCoV-229E    KTLQELSYKLPNYTVPDL..VVEQYNQTILNLTSEISTLENKSAELNYTVQKLQTLIDNI
HCoV-OC43    EELDQWFKNQTS.VAPDLSL..DYINVTFLDL..............QVEMNRLQEAIKVL
HCoV-NL63    KTLQEFAQNLPKYVKPNF..DLTPFNLTYLNLSSELKQLEAKTASLFQTTVELQGLIDQI
HCoV-HKU1    SELSHWFKNQTS.IAPNLTLNLHTINATFLDL..............YYEMNLIQESIKSL

                     1200      1210      1220
S-CryoEM     NESLIDLQELGKYEQYIKWPSGRLVPRGSPGSGYIPEAP.....................
SARS-CoV-2   NESLIDLQELGKYEQYIKWPWYIWLGFIA...GLIAIVMVTIMLCCMTSCCSCLK.....
SARS-CoV     NESLIDLQELGKYEQYIKWPWYVWLGFIA...GLIAIVMVTILLCCMTSCCSCLK.....
MERS-CoV     NESYIDLKELGNYTYYNKWPWYIWLGFIA...GLVALALCVFFILCCTGCGTNCM.....
HCoV-229E    NSTLVDLKWLNRVETYIKWPWWVWLCISV...VLIFVVSMLLLCCCSTGCCGFFSCFASS
HCoV-OC43    NHSYINLKDIGTYEYYVKWPWYVWLLICL...AGVAMLVLLFFICCCTGCGTSCF.....
HCoV-NL63    NSTYVDLKLLNRFENYIKWPWWVWLIISV...VFVVLLSLLVFCCLSTGCCGCCNCLTSS
HCoV-HKU1    NNSYINLKDIGTYEMYVKWPWYVWLLISF...SFIIFLVLLFFICCCTGCGSACF.....

                1230      1240      1250
S-CryoEM     .....RDGQ..AYV.RKDGEWVLLSTFLGHHHHHH
SARS-CoV-2   ..GCCSCGSCCKFD.EDDSEPVLKGVKLHYT....
SARS-CoV     ..GACSCGSCCKFD.EDDSEPVLKGVKLHYT....
MERS-CoV     ..GKLKCNRCCDRYEEYDLEPHKVHVH........
HCoV-229E    IRGCCE..STKLPYYDVEK......IHIQ......
HCoV-OC43    ..K..KCGGCCDDYTGYQE.LVIKTSHDD......
HCoV-NL63    MRGCCDCGSTKLPYYEFEK......VHVQ......
HCoV-HKU1    ..S..KCHNCCDEYGGHHD.FVIKTSHDD......
```

```
                          260       270       280       290
S-CryoEM     ..............G....DSSSGWTAGAAAYYVGYLQPRTFLLKYNENGTITDAVDCAL
SARS-CoV-2   ..............G....DSSSGWTAGAAAYYVGYLQPRTFLLKYNENGTITDAVDCAL
SARS-CoV     ...................PAQDIWGTSAAAYFVGYLKPTTFMLKYDENGTITDAVDCSQ
MERS-CoV     ..............Q....SDRKAW....AAFYVYKLQPLTFLLDFSVDGYIRRAIDCGF
HCoV-229E    TGHFYINGYRYFTLGNVEAVNFNVTTAETTDFCTVALASYADVLVNVSQTSIANIIYCN.
HCoV-OC43    ......................RRDIGFTLEYWVTPLTSRQYLLAFNQDGIIFNAVDCMS
HCoV-NL63    TGQFYINGFKYFDLGFIEAVNFNVTTASATDFWTVAFATFVDVLVNVSATNIQNLLYCD.
HCoV-HKU1    .....................SSNTDNETLQYWVTPLSKRQYLLKFDNRGVITNAVDCSS

               300       310       320       330       340       350
S-CryoEM     DPLSETKCTLKSFTVEKGIYQTSNFRVQPTESIVRFPNITNLCPFGEVFNATRFASVYAW
SARS-CoV-2   DPLSETKCTLKSFTVEKGIYQTSNFRVQPTESIVRFPNITNLCPFGEVFNATRFASVYAW
SARS-CoV     NPLAELKCSVKSFEIDKGIYQTSNFRVVPSGDVVRFPNITNLCPFGEVFNATKFPSVYAW
MERS-CoV     NDLSQLHCSYESFDVESGVYSVSSFEAKPSGSVVEQAEG.VECDFSPLLSG.TPPQVYNF
HCoV-229E    SVINRLRCDQLSFDVPDGFYSTSPIQSVELPVSIVSLPV....................Y
HCoV-OC43    DFMSEIKCKTQSIAPPTGVYELNGYTVQPIADVYRRKPDLPNCNIEAWLNDKSVPSPLNW
HCoV-NL63    SPFEKLQCEHLQFGLQDGFYSANFLDDNVLPETYVALPI....................Y
HCoV-HKU1    SFFSEIQCKTKSLLPNTGVYDLSGFTVKPVATVHRRIPDLPDCDIDKWLNNFNVPSPLNW

               360       370       380       390            400
S-CryoEM     NRKRISNCVADYSVLYNSASFSTFKCYGVSPTKLNDLCFTNV......YADSFVIRGDEV
SARS-CoV-2   NRKRISNCVADYSVLYNSASFSTFKCYGVSPTKLNDLCFTNV......YADSFVIRGDEV
SARS-CoV     ERKKISNCVADYSVLYNSTFFSTFKCYGVSATKLNDLCFSNV......YADSFVVKGDDV
MERS-CoV     KRLVFTNCNYNLTKLLSLFSVNDFTCSQISPAAIASNCYSSL......ILDYFSYPLSMK
HCoV-229E    HKHTFIVLY.................VDFKPQSGGGKCFNCYPAGVNITLANFNET...K
HCoV-OC43    ERKTFSNCNFNMSSLMSFIQADSFTCNNIDAAKIYGMCFSSI......TIDKFAIPNGRK
HCoV-NL63    YQHTDINFT....................ATASFGGSCYVCKPHQVNISLNG.......N
HCoV-HKU1    ERKIFSNCNFNLSTLLRLVHTDSFSCNNFDESKIYGSCFKSI......VLDKFAIPNSRR

             410       420       430       440             450
S-CryoEM     RQIAPGQTGKIADYNYKLPDDFTGCVIAWNSNNLDSKVGGN......YNYLYRLFR....
SARS-CoV-2   RQIAPGQTGKIADYNYKLPDDFTGCVIAWNSNNLDSKVGGN......YNYLYRLFR....
SARS-CoV     RQIAPGQTGVIADYNYKLPDDFMGCVLAWNTRNIDATSTGN......YNYKYRYLR....
MERS-CoV     SDLSVSSAGPISQFNYKQSFSNPTCLILATVPHNLTTITKP......LKYSYINKC....
HCoV-229E    GPLCVDTSHFTTKY.................VAVYA.....NVGRWSASINTGN.CPF..
HCoV-OC43    VDLQLGNLGYLQSSNYRIDTTATSCQLYYNLPAANVSVSRFNPSTWNKRFGFIEDSVFVP
HCoV-NL63    TSVCVRTSHFSIRY.................IYNRVKSGSPGDSSWHIYLKSGT.CPF..
HCoV-HKU1    SDLQLGSSGFLQSSNYKIDTTSSSCQLYYSLPAINVTINNYNPSSWNRRYGFNNFN....

                                       460             470       480
S-CryoEM     .........................KSNLKPFERDI.......STEIYQAGSTPCNGVEG
SARS-CoV-2   .........................KSNLKPFERDI.......STEIYQAGSTPCNGVEG
SARS-CoV     .........................HGKLRPFERDI.......SNVPFSPDGKPCTP.PA
MERS-CoV     .........................SRLLSDDRTEV.......PQLVNANQYSPCVSIVP
HCoV-229E    .SFGK.VNNFVKFGSVCFSLKDIPGGCAM.............................PI
HCoV-OC43    QPTGVFTNHSVVYAQHCFKAPKNFCPCKL...NGSCPGKNNGIGTCPAGTNYLTCDNLC.
HCoV-NL63    .SFSK.LNNFQKFKTICFSTVEVPGSCNF.............................PL
HCoV-HKU1    .....LSSHSVVYSRYCFSVNNTFCPCAKPSFASSCKSHKPPSASCPIGTNYRSCESTT.

                         490                         500       510
S-CryoEM     FNC..........YF..PLQSYGFQP.............TNGVGYQ..PYRVVV......
SARS-CoV-2   FNC..........YF..PLQSYGFQP.............TNGVGYQ..PYRVVV......
SARS-CoV     LNC..........YW..PLNDYGFYT.............TTGIGYQ..PYRVVV......
MERS-CoV     STVWEDGDYYRKQLS..PLEGGGWLV.............ASGSTVA..MTEQLQ......
HCoV-229E    VANWAYSKYYT.................................................
HCoV-OC43    ..TLDPI..........TFKAPGTYKCPQTKSLVGIGEHCSGLAVKSDYCR......GNS
HCoV-NL63    EATWHYTSYTI.................................................
HCoV-HKU1    ..VLDHTDWCRCSCLPDPITAYDPRSCSQKKSLVGVGEHCAGFGVDEEKCGVLDGSYNVS

                                         520            530       540
S-CryoEM     ........LS..............FELL....HAPATVCGP.....KKSTNLVKNKCVNF
SARS-CoV-2   ........LS..............FELL....HAPATVCGP.....KKSTNLVKNKCVNF
SARS-CoV     ........LS..............FELL....NAPATVCGP.....KLSTDLIKNQCVNF
MERS-CoV     ........MG..............FGITVQYGTDTNSVCPKLEFANDTKIASQLGNCVEY
HCoV-229E    ...IGSLYVSWSDGDGITGVP..............QPVE.....GVSSFMNVTLDKCTKY
HCoV-OC43    CTCQPQAFLGWSADSCLQGDKCNIFANLILHDVNSGLTCST..DLQKANTDIILGVCVNY
HCoV-NL63    ...VGALYVTWSEGNSITGVP..............YPVS.....GIREFSNLVLNNCTKY
HCoV-HKU1    CLCSTDAFLGWSYDTCVSNNRCNIFSNFILNGINSGTTCSN..DLLQPNTEVFTDVCVDY
```

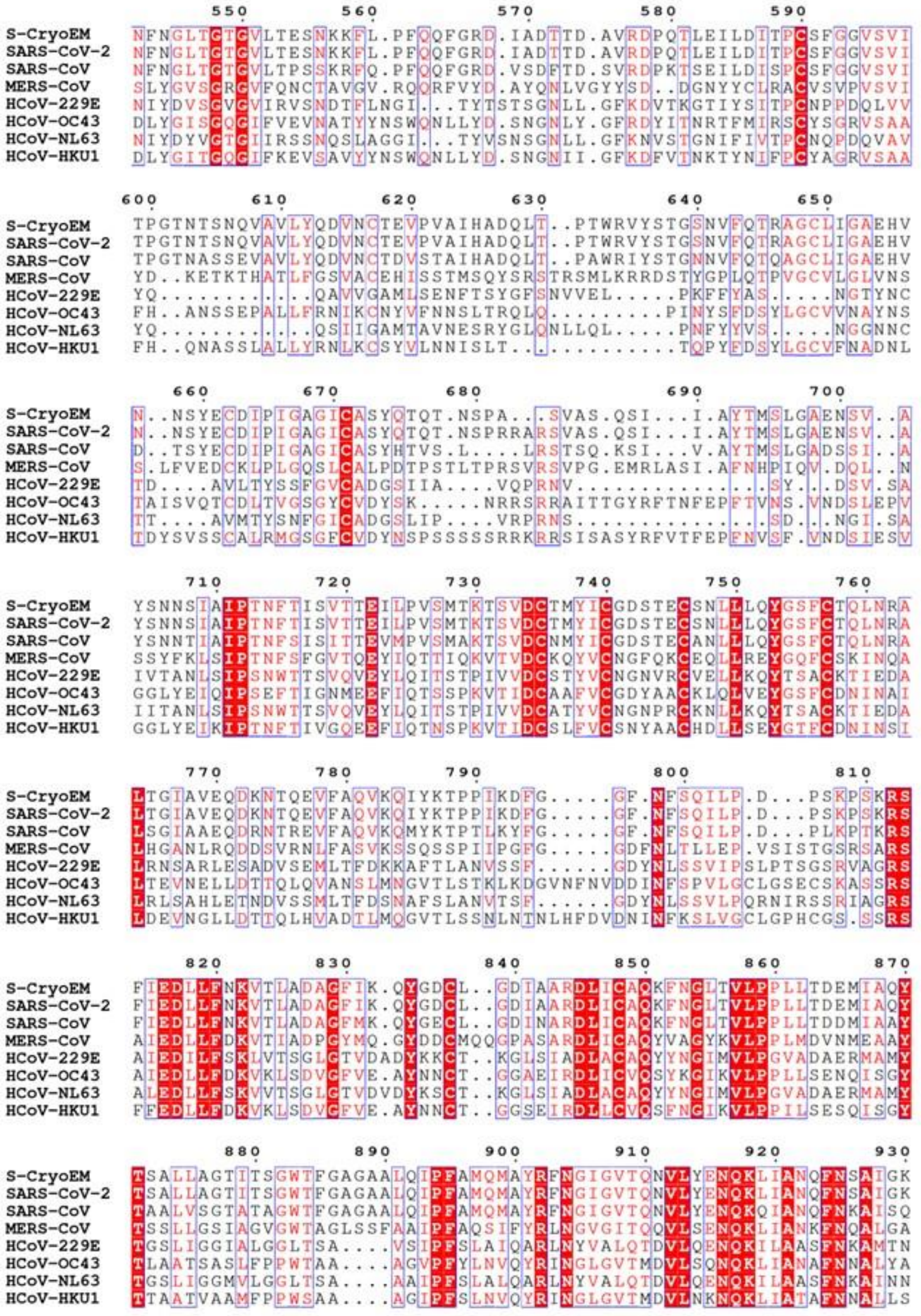

```
                                                940       950       960       970
S-CryoEM    IQDSLSSTA..............SALGKLQDVVNQNAQALNTLVKQLSSNFGAISSVLND
SARS-CoV-2  IQDSLSSTA..............SALGKLQDVVNQNAQALNTLVKQLSSNFGAISSVLND
SARS-CoV    IQESLTTTS..............TALGKLQDVVNQNAQALNTLVKQLSSNFGAISSVLND
MERS-CoV    MQTGFTTTN..............EAFRKVQDAVNNNAQALSKLASELSNTFGAISASIGD
HCoV-229E   IVDAFTGVNDAITQTSQALQTVATALNKIQDVVNQQGNSLNHLTSQLRQNFQAISSSIQA
HCoV-OC43   IQQGFDATN..............SALVKIQAVVNANAEALNNLLQQLSNRFGAISASLQE
HCoV-NL63   IVASFSSVNDAITQTAEAIHTVTIALNKIQDVVNQQGSALNHLTSQLRHNFQAISNSIQA
HCoV-HKU1   IQNGFSATN..............SALAKIQSVVNSNAQALNSLLQQLFNKFGAISSSLQE

              980       990      1000      1010      1020      1030
S-CryoEM    ILSRLDKVEAEVQIDRLITGRLQSLQTYVTQQLIRAAEIRASANLAATKMSECVLGQSKR
SARS-CoV-2  ILSRLDKVEAEVQIDRLITGRLQSLQTYVTQQLIRAAEIRASANLAATKMSECVLGQSKR
SARS-CoV    ILSRLDKVEAEVQIDRLITGRLQSLQTYVTQQLIRAAEIRASANLAATKMSECVLGQSKR
MERS-CoV    IIQRLDVLEQDAQIDRLINGRLTTLNAFVAQQLVRSESAALSAQLAKDKVNECVKAQSKR
HCoV-229E   IYDRLDTIQADQQVDRLITGRLAALNVFVSHTLTKYTEVRASRQLAQQKVNECVKSQSKR
HCoV-OC43   ILSRLDALEAEAQIDRLINGRLTALNAYVSQQLSDSTLVKFSAAQAMEKVNECVKSQSSR
HCoV-NL63   IYDRLDSIQADQQVDRLITGRLAALNAFVSQVLNKYTEVRGSRRLAQQKINECVKSQSNR
HCoV-HKU1   ILSRLDALEAQVQIDRLINGRLTALNAYVSQQLSDISLVKFGAALAMEKVNECVKSQSPR

             1040      1050      1060      1070      1080      1090
S-CryoEM    VDFCGKGYHLMSFPQSAPHGVVFLHVTYVPAQEKNFTTAPAICHDGKA...HFPREGVFV
SARS-CoV-2  VDFCGKGYHLMSFPQSAPHGVVFLHVTYVPAQEKNFTTAPAICHDGKA...HFPREGVFV
SARS-CoV    VDFCGKGYHLMSFPQAAPHGVVFLHVTYVPSQERNFTTAPAICHEGKA...YFPREGVFV
MERS-CoV    SGFCGQGTHIVSFVVNAPNGLYFMHVGYYPSNHIEVVSAYGLCDAANPTNCIAPVNGYFI
HCoV-229E   YGFCGNGTHIFSIVNAAPEGLVFLHTVLLPTQYKDVEAWSGLCVDGTN..GYVLRQPNLA
HCoV-OC43   INFCGNGNHIISLVQNAPYGLYFIHFNYVPTKYVTAKVSPGLCIAGNR..GIAPKSGYFV
HCoV-NL63   YGFCGNGTHIFSIVNSAPDGLLFLHTVLLPTDYKNVKAWSGICVDGIY..GYVLRQPNLV
HCoV-HKU1   INFCGNGNHILSLVQNAPYGLLFMHFSYKPISFKTVLVSPGLCISGDV..GIAPKQGYFI

                   1100      1110      1120      1130          1140
S-CryoEM    SN.....GTHWFVTQRNFYEPQIITTDNTFVSGNCDVVIGIVNNTVYDPLQ..PELDSFK
SARS-CoV-2  SN.....GTHWFVTQRNFYEPQIITTDNTFVSGNCDVVIGIVNNTVYDPLQ..PELDSFK
SARS-CoV    FN.....GTSWFITQRNFFSPQIITTDNTFVSGNCDVVIGIINNTVYDPLQ..PELDSFK
MERS-CoV    KTNNTRIVDEWSYTGSSFYAPEPITSLNTKYVAPQVTYQN.ISTNLPPPLLGNSTGIDFQ
HCoV-229E   LY...KEGNYYRITSRIMFEPRIPTMADFVQIENCNVTFVNISRSEL..QTIVPEYIDVN
HCoV-OC43   NV.....NNTWMYTGSGYYYPEPITENNVVVMSTCAVNYTKAPYVML..NTSIPNLPDFK
HCoV-NL63   LY...SDNGVFRVTSRVMFQPRLPVLSDFVQIYNCNVTFVNISRVEL..HTVIPDYVDVN
HCoV-HKU1   KH.....NDHWMFTGSSYYYPEPISDKNVVFMNTCSVNFTKAPLVYL..NHSVPKLSDFE

             1150      1160      1170                     1180      1190
S-CryoEM    EELDKYFKNHTS.PDVDLG.DISGINASVVNI..............QKEIDRLNEVAKNL
SARS-CoV-2  EELDKYFKNHTS.PDVDLG.DISGINASVVNI..............QKEIDRLNEVAKNL
SARS-CoV    EELDKYFKNHTS.PDVDLG.DISGINASVVNI..............QKEIDRLNEVAKNL
MERS-CoV    DELDEFFKNVST.SIPNFG.SLTQINTTLLDL..............TYEMLSLQQVVKAL
HCoV-229E   KTLQELSYKLPNYTVPDL..VVEQYNQTILNLTSEISTLENKSAELNYTVQKLQTLIDNI
HCoV-OC43   EELDQWFKNQTS.VAPDLSL..DYINVTFLDL..............QVEMNRLQEAIKVL
HCoV-NL63   KTLQEFAQNLPKYVKPNF..DLTPFNLTYLNLSSELKQLEAKTASLFQTTVELQGLIDQI
HCoV-HKU1   SELSHWFKNQTS.IAPNLTLNLHTINATFLDL..............YYEMNLIQESIKSL

                  1200      1210      1220
S-CryoEM    NESLIDLQELGKYEQYIKWPSGRLVPRGSPGSGYIPEAP.....................
SARS-CoV-2  NESLIDLQELGKYEQYIKWPWYIWLGFIA...GLIAIVMVTIMLCCMTSCCSCLK.....
SARS-CoV    NESLIDLQELGKYEQYIKWPWYVWLGFIA...GLIAIVMVTILLCCMTSCCSCLK.....
MERS-CoV    NESYIDLKELGNYTYYNKWPWYIWLGFIA...GLVALALCVFFILCCTGCGTNCM.....
HCoV-229E   NSTLVDLKWLNRVETYIKWPWWVWLCISV...VLIFVVSMLLLCCCSTGCCGFFSCFASS
HCoV-OC43   NHSYINLKDIGTYEYYVKWPWYVWLLICL...AGVAMLVLLFFICCCTGCGTSCF.....
HCoV-NL63   NSTYVDLKLLNRFENYIKWPWWVWLIISV...VFVVLLSLLVFCCLSTGCCGCCNCLTSS
HCoV-HKU1   NNSYINLKDIGTYEMYVKWPWYVWLLISF...SFIIFLVLLFFICCCTGCGSACF.....

               1230      1240      1250
S-CryoEM    .....RDGQ..AYV.RKDGEWVLLSTFLGHHHHHH
SARS-CoV-2  ..GCCSCGSCCKFD.EDDSEPVLKGVKLHYT....
SARS-CoV    ..GACSCGSCCKFD.EDDSEPVLKGVKLHYT....
MERS-CoV    ..GKLKCNRCCDRYEEYDLEPHKVHVH........
HCoV-229E   IRGCCE..STKLPYYDVEK......IHIQ......
HCoV-OC43   ..K..KCGGCCDDYTGYQE.LVIKTSHDD......
HCoV-NL63   MRGCCDCGSTKLPYYEFEK......VHVQ......
HCoV-HKU1   ..S..KCHNCCDEYGGHHD.FVIKTSHDD......
```
